# Gaze Following in Pigeons Increases with the Number of Demonstrators

**DOI:** 10.1101/2024.12.05.627032

**Authors:** Mathilde Delacoux, Akihiro Itahara, Fumihiro Kano

## Abstract

Gaze following, orienting one’s gaze in the same direction as another individual, is widespread across species and well-studied in terms of cognitive mechanisms. However, the role of collective context on this behavior is less understood. This study tested whether the number of conspecifics looking in a particular direction facilitates gaze following in pigeons (*Columba livia*). We presented attention-getting stimuli to subsets of pigeons (demonstrators), while other pigeons (observers) could not see the stimuli directly. We used a motion-capture system to track fine-scale movements of head and body orientations. Observer pigeons followed the gaze of demonstrators, specifically toward the target object rather than a perceptually similar distractor. Critically, the likelihood and frequency of gaze following increased with the number of demonstrators looking at the target. We observed no nonlinear effects, such as saturation or quorum thresholds, in this increasing trend, at least under our experimental conditions. In group-living species like pigeons, multiple individuals looking in the same direction may serve as a more reliable social signal. To our knowledge, this is the first study to demonstrate that the number of demonstrators influences gaze following in non- human animals, highlighting the critical role of collective context in animal social cognition.

## Introduction

In many socially living animals, individual cognition is embedded within the group context and shaped by interactions among multiple individuals. Previous studies on animal collective behavior have emphasized its importance, as demonstrated by collective perception in fish schools (Berdahl et al., 2013), consensus-driven decision-making by ants, fishes (Couzin, 2009; Sumpter, 2006) and baboons (Strandburg-Peshkin et al., 2015), and cultural transmission of social information in bumblebees (Bridges et al., 2024), great tits (Aplin et al., 2015; Chimento et al., 2021) and great apes (Whiten et al., 2007). Despite these advancements, previous studies on animal social cognition has primarily been constrained to experimental settings with only a limited number of animals, typically dyads, rather than multiple individuals. Although the dyadic setup allows for controlled observation of both the behavior of stimulus animals and the responses of observer animals—making it particularly useful for studying the cognitive mechanisms behind individual socio-cognitive behaviors (Balda et al., 1998; Heyes & Jr, 1996)—it overlooks key aspects of social complexity found in real-world settings.

Gaze following, defined as orienting one’s gaze in the same direction as another individual, presents an interesting research topic for exploring the effect of collective contexts on social cognition. This behavior is one of the most extensively studied socio-cognitive behaviors in animal cognition research, particularly from the perspectives of development, phylogeny, and cognitive mechanisms (Davidson et al., 2014; Del Bianco et al., 2019; Zeiträg, 2022; Zuberbühler, 2008). Research has shown that gaze following is widespread across various taxa, including birds (Bugnyar et al., 2004; Butler & Fernández-Juricic, 2014; Kehmeier et al., 2011; Loretto et al., 2010; Nawroth et al., 2017; Zeiträg et al., 2023), primates (Bräuer et al., 2005; Kano & Call, 2014; Shepherd & Platt, 2008; Tomasello et al., 1998), non-primate mammals (Kaminski et al., 2005; Met et al., 2014), reptiles (Wilkinson et al., 2010; Zeiträg et al., 2023), and likely even some fish (Goossens, 2008; Leadner et al., 2021). While basic co-orienting of gaze may result from reflexive behavior copying, more complex pattern of behaviors, such as gaze following around barriers, likely indicate more complex cognitive processes, including perspective taking (Davidson et al., 2014; Davidson & Clayton, 2016; Tomasello et al., 1999; Zeiträg, 2022; Zuberbühler, 2008). The gaze following response is also modulated by several contextual factors, such as dominance

(Shepherd et al., 2006), sex (Deaner et al., 2006), whether the demonstrator is a conspecific or allospecific (Hattori et al., 2010; Kano & Call, 2014), facial expression (Gallup et al., 2014; Goossens et al., 2008), or communicative cues like eye contact (Senju & Csibra, 2008; Téglás et al., 2012; but see Gredebäck et al., 2018; Kano et al., 2018). These varying levels of response are thought to engage different neural pathways: one is the subcortical route, which is fast and reflexive, while the other is the cortical pathway, which integrates social and contextual information to understand gaze in a more context-dependent manner (Shepherd, 2010). While basic co-orienting behavior is observed early in life, gaze following around barriers as a form of perspective-taking, typically develops later in both human and non- human animals (Farroni et al., 2004; Moll & Tomasello, 2004; Scaife & Bruner, 1975; Schloegl et al., 2007).

Despite the extensive research focus on its cognitive mechanisms, little is known about how gaze following occurs in a group, beyond a dyad. In humans, Milgram et al. (1969) pioneered a field experiment where a varying number of confederate demonstrators looked up at the top of a building on a street. They found that the larger the number of demonstrators exhibiting a gaze cue, the more likely passersby were to follow their gaze. Although Milgram et al. suspected that this response was quorum-like – a response triggered only after a critical threshold is reached, a key feature often observed in collectives (Sumpter & Pratt, 2009) – a more recent follow-up study by Gallup et al. (2012) found no support for this claim. In a similar experiment, they found that the passersby’s response followed a proportional saturating relationship, meaning that an initial increase in the number of demonstrators led to a corresponding proportional increase in the observers’ looking response, but then the response gradually saturated and reached a plateau.

To our knowledge, no study has examined the effect of the number of demonstrators on gaze following in nonhuman animals. This study investigated the gaze-following responses of pigeons, where a varying number of demonstrators provided gaze cues to observers within a flock. Pigeons are well-suited to this task because they typically form relatively large flocks, where individuals are usually surrounded by multiple conspecifics (both in captivity and in the wild). In such flocks, following another’s gaze should be beneficial for key activities such as foraging and vigilance. The gaze cues provided by one versus multiple individuals might convey different meanings to an individual, as locations attended by a larger number of individuals potentially indicate the presence of more significant information.

We used a recently developed motion-capture-based posture tracking system for a flock of pigeons, where all individuals’ head and body orientations were tracked with high spatiotemporal precision, allowing them to naturally flock together in an unconstrained space and engage in activities such as foraging and vigilance (Delacoux & Kano, 2024; Kano et al., 2022; Nagy et al., 2023). This setup, when combined with a typical gaze-following experiment, is expected to capture any subtle cues provided by demonstrator animals and gaze-following responses from observer animals.

Although there is no direct evidence of gaze following in pigeons, several species of birds have been shown to follow the gaze of a conspecific or a human experimenter (e.g., ravens: Bugnyar et al., 2004; starlings: Butler & Fernández-Juricic, 2014; geese: Kehmeier et al., 2011; ibises: Loretto et al., 2010; pinguins: Nawroth et al., 2017; tinamous, emus, rheas and jungle fowls: Zeiträg et al., 2023). We thus expected that pigeons also follow the gaze of at least their conspecifics.

This study aimed to answer four questions: 1) whether pigeons exhibit gaze-following responses, 2) whether a larger number of demonstrators enhance the observers’ gaze- following responses, and 3) if so, whether the increase in response follows a linear, quorum, or saturation pattern. As a secondary question, we also asked 4) whether pigeons could distinguish the gaze target of the demonstrators from a distractor object (a perceptually equivalent object). This last question has not been addressed in previous studies on bird gaze following, largely due to the lack of a fine-scale tracking system. This question is particularly interesting because many bird species have two or more types of ‘gaze’; in pigeons, there are two laterally projecting foveas, one area projecting into the lower-frontal visual field for close-range foraging (the projection of the ‘red field’), and another projecting more frontally into the binocular field, which is often used for pecking (Kano et al., 2022), perching (Green et al., 1992) or attending to slowly moving objects (Maldonado et al., 1988). Thus, it remains unclear whether observer pigeons can identify the gaze target of demonstrators when both target and distractor objects are present, despite the multiple possibilities for gaze direction.

## Methods

### Subjects

A total of 70 pigeons were used in this study (24 in the first experiment and 46 in the second). They were housed in an outdoor aviary 2x4x2m at the Max-Planck Institute of Animal Behavior, Radolfzell, Germany. They had *ad libidum* access to water and grit and were fed once a day with a mix of seeds.

### Ethics statement

Animal experiments in this study were performed under the licenses 35-9185.81/G-19/107 and 35-9185.81/G-22/100 granted by the Regierungspräsidium Freiburg, Abteilung Landwirtschaft, Ländlicher Raum, Veterinär- und Lebensmittelwesen, animal ethics authorities of Baden-Württemberg, Germany. No animal was injured or killed during the tests and handling was kept minimal to reduce stress. Outside of experiments, the pigeons were socially housed in their home aviary, provided with perches and nesting structures, and checked daily for health state. After the daily experiment was completed, the markers were removed from the pigeons’ heads and backpacks, and they were brought back to their loft.

### Experimental design

#### Rationales and basic design

This study consisted of 3 experiments aimed at demonstrating gaze following in pigeons by progressively refining the experimental setup throughout the experiments. Also, to ensure sufficient data for analysis, we combined the results of Experiments 1 and 3. Experiment 1 was performed with a flock of pigeons (n = 24) in a group setup. In the test condition, a varying number of pigeons from the flock of ten could see a moving object while the others could not see. In the control condition, none of the pigeons could see the moving object, thereby establishing a baseline observation as well as controlling for the potential effect of sound from the moving object. In Experiment 1, while we observed an effect of the number of demonstrators on the observer’s behavior, there was no significant difference between test and control conditions. Experiment 2 tested the same pigeons and aimed to confirm the gaze-following responses in a more conventional setting, in a dyadic setup with either a conspecific, or a human as a demonstrator (respectively Exp 2 and Exp 2 follow-up: more details in Suppl. Mat. 1). In addition, we modified the control condition (see below). Although we found some indication of gaze following in this experiment, the difference between test and control conditions did not reach statistical significance. Given that we expect a collective effect on gaze following, Experiment 3 tested a larger number of new pigeons (n = 46), while performing fewer trial for each individual to avoid habituation of the demonstrators, and adopting the improved control design.

### Set-ups

#### Motion-capture system

The experiments were conducted in the SMART-BARN (Nagy et al., 2023; Fig 1a). The setup included a motion-capture system with 32 infrared cameras covering a 15x7x4 meter volume. At the center, an inverted U-shaped structure (3w x 2h m (w = width, h = height)) held 8 opaque tubes (4 on top, 4 on the ground, Fig 1b), each containing a small colorful object (4-5 cm diameter) attached to a string control system. Each tube had a small window from which the object is visible only from one side Fig 1c. A set of pipes was leading the strings coming out of the tube structure to a control station, hidden from the pigeons’ sight by a curtain, where the experimenter can control the objects movements. Pigeons were placed on two wooden tables (210l x 22w x 75h cm; l = length) positioned parallel to the structure, where they could either see (“demonstrators”) or could not see (“observers”) the moving objects through the window. During experiments, the objects were moved back and forth in either the top or bottom tubes. The location presenting the objects is referred as the “target” location while the other location not presenting the objects is referred as the “distractor” location. After habituation to the structure and tables, pigeons naturally preferred standing on the tables and no particular training was required.

**Figure 1:**
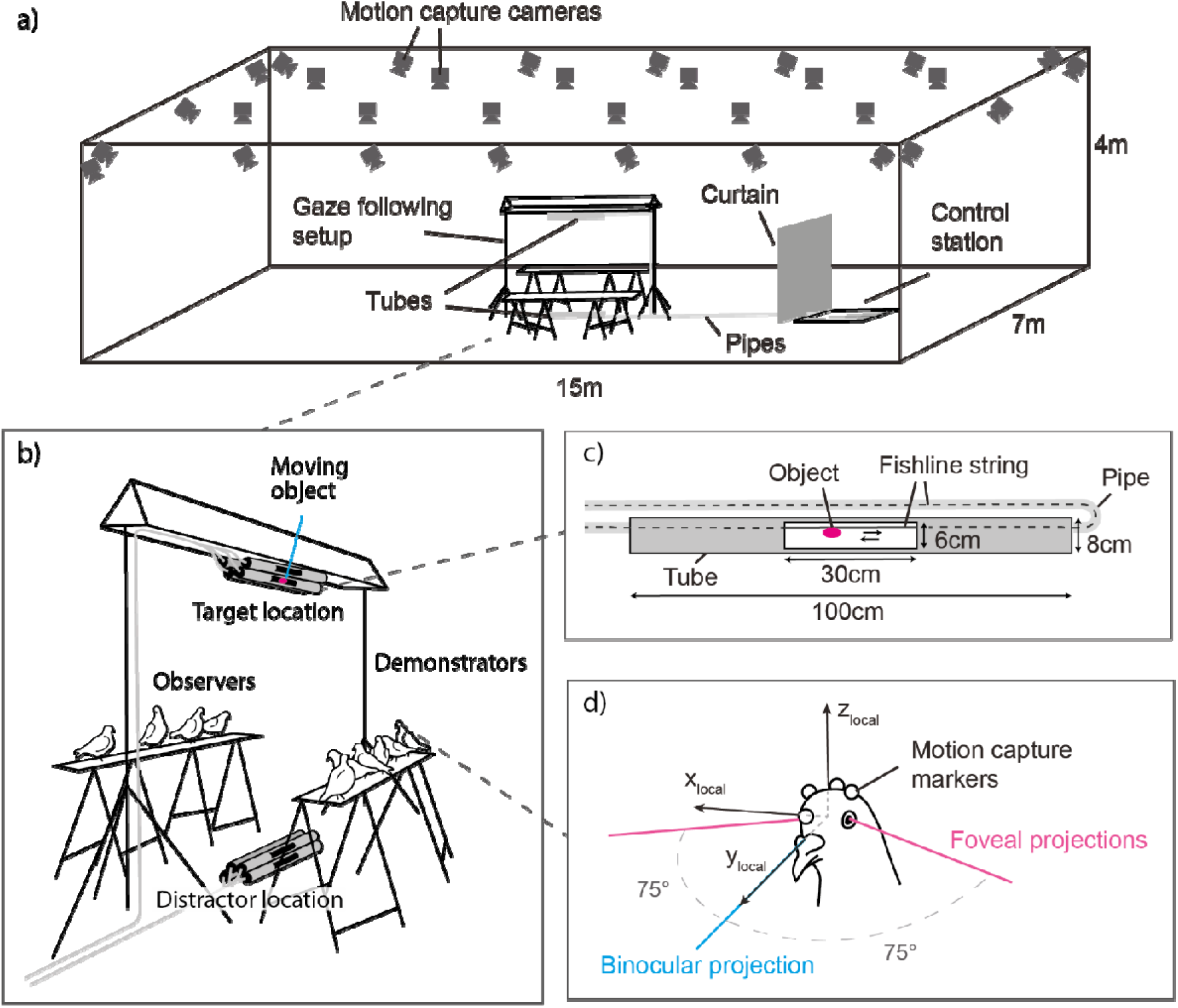
Experimental set-up. a) The experiment took place in the SMART-BARN tracking system developed by the Max Planck Institute of Animal Behavior (Nagy et al., 2023), with the experimental setup located at its center. b) Ten pigeons are released on the tables and can be arranged on both sides of the tubes structure by the experimenter to obtain various numbers of demonstrators and observers. Two sets of tubes were placed at the bottom and top of the structure, one containing a moving object (target location) and one not displaying anything (distractor location) to the demonstrators. c) Small colorful objects attached to a fishline string system are presented from inside a tube, from which a window is cut, making the object visible from one side only. d) Using the 4 motion capture markers attached to the pigeon’s head, we calculated the foveal and binocular projections in the head local coordinate system.

Prior to the daily experiment, the motion capture cameras were calibrated using a standard procedure with Active Wand (VICON; calibration with 2000 frames per camera).

### Marker attachment and eye-beak calibration

Four motion capture markers (6.4mm diameter, OptiTrack) were placed on the head of the pigeons and a Styrofoam plate assorted with a unique combination of 4 markers (9mm diameter) for identification was attached to their back. The head of the pigeons was recorded from 4 different angles with synchronized webcams. This procedure (further detailed in Kano et al., 2022; Nagy et al., 2023) allowed for the reconstruction of the eyes and beak tip positions relative to the head markers.

### Procedures

#### General

Each pigeon experienced no more than a trial per day. In the experiment, the pigeons were released on tables on either side of the structure. Within each trial, they were exposed to a succession of object presentations. In each object presentation, the experimenter, hidden behind a curtain, pulled fish lines attached to the object back-and-forth twice for about 24 seconds. Before and after each object presentation, the object was placed in the occluded sections of the tubes. The interval between object presentations was varied randomly (min 20 seconds, mean 54 seconds), to prevent learning of the object appearances by the pigeons.

#### Group set-up (Exp 1 and 3)

Ten pigeons were released on the tables. The pigeons’ locations were shuffled between presentation by the experimenter gently approaching one of the tables, triggering flight responses to the other table, ensuring that all pigeons experienced different numbers of demonstrators. In the test condition, identical in both Exp. 1 and 3, demonstrators saw the moving object and could provide gaze cues to observers.

However, Exp. 1 and Exp. 3 differ in the control condition and in the number of trials and object presentations. In the control condition of Exp. 1, all pigeons were moved to one side (observers only), and objects were presented on the opposite side (no demonstrator; Suppl. Fig S1). In the control condition of Exp. 3 (as well as Exp. 2), the object was presented in an occluded tube (no window) while keeping the numbers and identities of demonstrator and observer pigeons as similar as possible between conditions (Suppl. Fig S1). By performing the test and control conditions in two successive trials, this allowed for a better pairwise comparison. For the number of trials, Exp. 1 had 8 test and 4 control object presentations (we required more test presentations than controls, as only observers provided data in tests, while all 10 pigeons contributed data in controls) in pseudo-counterbalanced order across trials (either TTTTCCTTTTCC or CCTTTTCCTTTT; with T = one test presentation and C = one control presentation), and each pigeon underwent 4-6 trials. Exp. 3 had 8 test and 8 control object presentations (only pigeons standing on the “observer” table contributed to both the test and control data) in a randomly counterbalanced order across trials (e.g., TC- CT-TC-TC-CT-TC-CT-CT), and each pigeon underwent 2-3 trials (fewer trials to avoid habituation).

#### Dyadic set-up (Exp 2)

In Experiment 2, we presented objects only on the top tubes (no bottom tube). Two pigeons were placed on opposite sides of a small table and separated by a thin net. Each trial consisted of two test presentations (with each pigeon acting once as a demonstrator and once as an observer) and one control presentation, where the object was in an occluded tube and no pigeon could see it. The sequence was randomized, and each pigeon experienced six trials on different days with different pairs (one trial per day). As a follow-up, we also performed an experiment with the identical design except that the demonstrator is a human experimenter as in some previous studies (ravens: Bugnyar et al., 2004; parrots: Giret et al., 2009; bonobos: Kano & Call, 2014; dogs: Met et al., 2014; jackdaws: Von Bayern & Emery, 2009) to present longer and more standardized gaze cues. An experimenter stood on one side and looked up during the test and looked behind the pigeon during the control (see Suppl. Mat. 1).

### Data analysis

#### Mo-cap data processing

The details of the pipelines used to process the motion capture data can be found in Delacoux & Kano (2024) and the code is available in Delacoux & Kano (2023). Briefly, the motion capture coordinates were exported as a CSV file using the Nexus software (version 2.14, VICON). The data was then screened by the pipeline to correct for mislabeling of the individuals’ ID and marker positions. The relative positions of eyes and beak to the attached head markers were reconstructed using a custom Python computer vision model. From the positions of eyes and beak, the head local coordinate system was reconstructed with the midpoint of the two eyes as the origin and the Y axis pointing to the horizon (defined as 30 degrees above the origin-beak line in elevation). The reconstructed head was then applied onto the 3D coordinates tracked by the motion capture system to define the head local coordinate system during the whole trial, then filtered and smoothed to eliminate noise.

#### Definition of looking

From the head local coordinate system, we reconstructed the visual field of the individuals. As in a previous studies (Delacoux & Kano, 2024), foveal projections were defined as visual cones with their center lines projecting ±75° in azimuth and 0° in elevation in the head’s local coordinate system, with 10° error margin to accommodate for eye movement and noise.

Consistent with the previous studies, our pigeons primarily used one of their foveas to look at the objects in this study. Interestingly, they occasionally also used their binocular field, consistent with the previous observation that they use binocular field when attending to slow-moving objects or when perching (Green et al., 1992; Maldonado et al., 1988). We observed no use of the “red field” in the pigeons’ eyes, typically used to attend to close ground objects in the previous studies. We included both foveal and binocular field use into the analysis, with the latter defined as a cone with its center line projecting 0° in both azimuth and elevation Fig 1d. We observed that pigeons used their foveas and binary visual fields to look at the moving object in 92% and 8% of the cases respectively.

The region of interest for target or distractor object was defined as a sphere of 50 cm diameter encompassing the window’s 30 cm width, with a 10 cm margin on each side.

Looking was defined when at least one of its reconstructed gaze cones crossed the spheres. Several filters were applied to reduce random gaze crossings. First, we excluded the frames when the individual was making head saccades, during which visual processing is likely inhibited in birds (Brooks & Holden, 1973), and also when the individual was grooming, as the individual was likely not attending to the environment. Grooming was automatically detected using a predefined algorithm (Delacoux & Kano, 2024). Second, we only considered fixations longer than 300 ms onto the object. Third, to ensure that only relevant demonstrator looks were considered for each observer, demonstrator looks were excluded when the observer was grooming.

### Data exclusion

Pigeons that flew off or switched tables during the presentation, and test presentations where no demonstrator looked at the target were excluded from the analysis (Exp. 1: 7 of 842 observations; Exp. 2: 8 of 288 observations; Exp. 3: 20 of 690 observations).

### Statistical analysis

We fitted the data with generalized linear mixed models (GLMMs). The main response variables were the likelihood of an observer looking at either the target or distractor location (in binomial models) and the number of looks at the target or distractor location per presentation (in Poisson models). The main predictor were the condition (test or control) to test the presence of gaze following in each experiment and the “actual number of demonstrators” to test the effect of a collective effect on gaze following. The latter variable represented the number of pigeons on the demonstrator side that gave a gaze cue, i.e. that looked at the target at least once during the presentation. For all models, we also included control predictors, i.e., the trial number, the presentation number, the presentation location (for Exp. 1 and 3, as Exp. 2 only had the top location), the experiment number (1 or 3, only for the combined analysis), the random effect of the individual ID, and random slopes for all fixed effects.

For all models, all numerical variables were normalized to control for scale effects.

Detailed R formulas are provided in Suppl. Table S1. In addition, we checked the collinearity of the explanatory variables in all models, and for Poisson models, we checked for overdispersion.

## Results

### Gaze following of pigeons

To test the presence of gaze following in pigeons, we tested whether the observers’ response (likelihood of looking and the number of looks at the target location) was predicted by the condition (test condition, where demonstrators see the moving object and give a gaze cue; and control condition, where there is either no demonstrators (Exp. 1) or the demonstrators do not see the moving object (Exp. 2 and 3). For both Exp. 1 and 2, we did not find a significant effect of the condition on neither the observers’ likelihood of looking (Exp. 1: χ2(1) = 0.113, β = -0.066, *p= 0.736*; Exp.2: χ2(1) = 1.848, β = 0.443, *p= 0.174*; Fig 2a, for more details about model results see Suppl. Table S2) nor the observers’ number of looks (Exp.1: χ2(1) = 0.077, β = 0.061, *p= 0.782*; Exp. 2: χ2(1) = 2.006, β = 0.335, *p= 0.157;* Fig 2b). In Exp. 3, the condition predicted both the likelihood of looking (χ2(1) = 4.964, β = 0.448, *p= 0.026*; Fig 2a) and the number of looks of the observers (χ2(1) = 6.574, β = 0.425, *p= 0.010*; Fig 2b); the pigeons were more likely to look at the target and looked more times in the test compared to the control condition.

**Figure 2:**
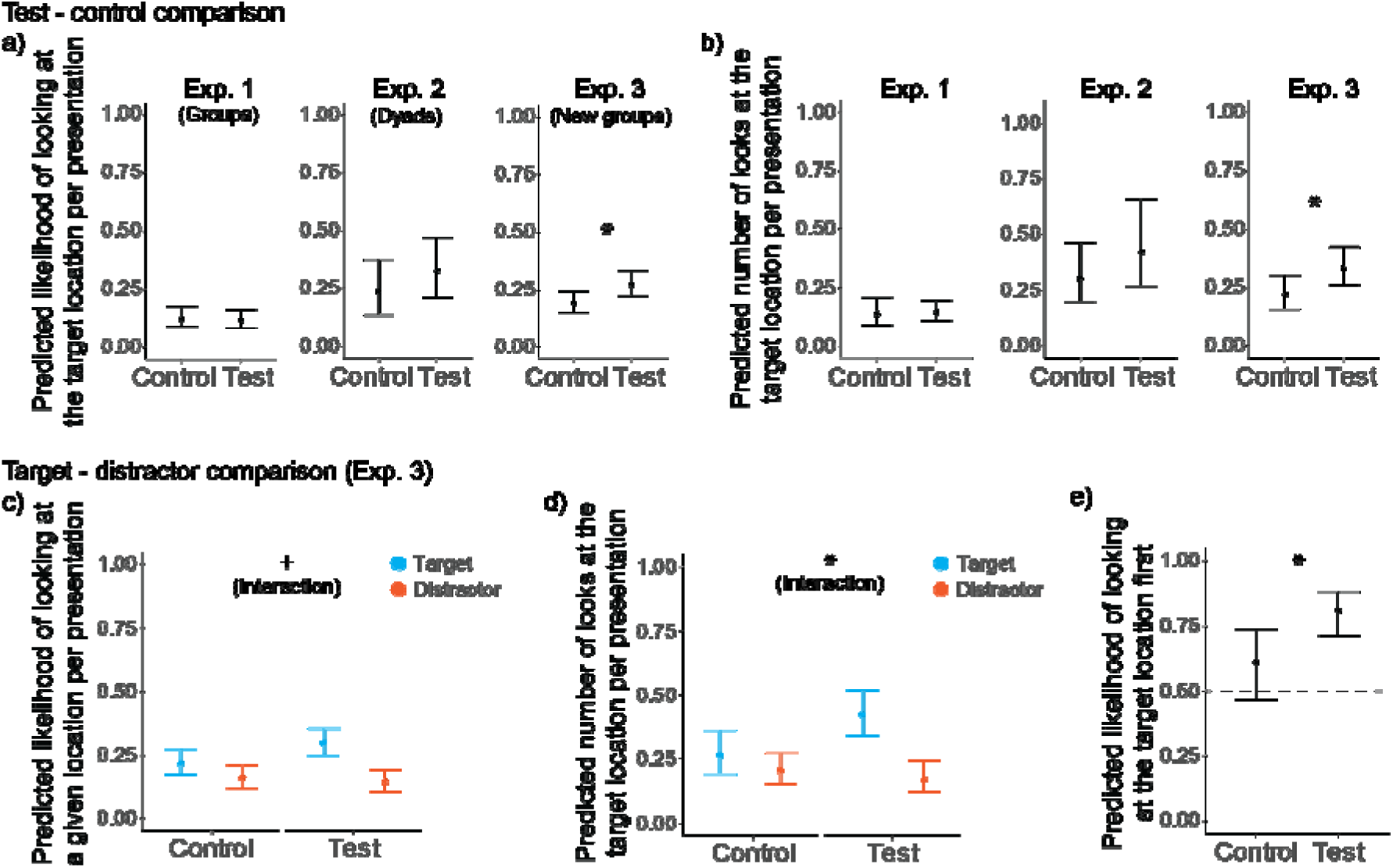
Gaze following results of pigeons. a-b) Test-Control comparison: standard way of testing gaze following in dyadic setups. The graphs show the looking response (a) likelihood of looking and b) number of looks) as a function of condition. c-e) Comparison of the response towards the target vs distractor location in the test and the control conditions in the Exp. 3, to test if the pigeons responded specifically more to the target compared to the distractor location in the test setup. The graphs show the looking response (c) likelihood of looking, d) number of looks and e) likelihood of looking at the target first) as a function of condition for the target and distractor locations. For c-d), symbols denotates the significance of the interaction condition*location. For all graphs, confidence intervals (CI) were calculated using 10 000 simulations of the model parameters and represents the range between the 2.5th and 97.5th percentiles of the simulated predictions. p= 0.1 > + > 0.05 > * > 0.01 > ** > 0.001 > ***

**Figure 3:**
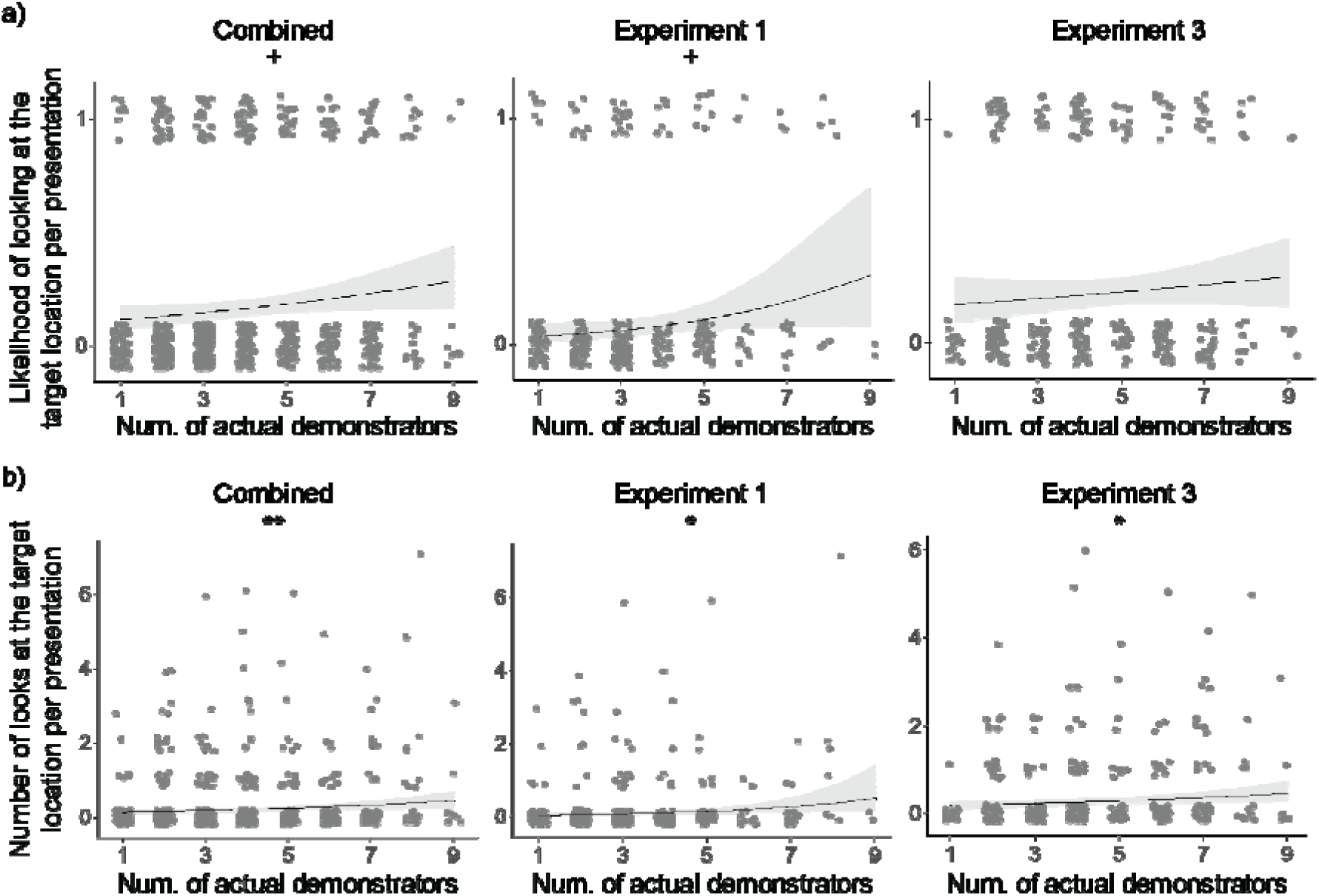
Looking response (a) likelihood of looking and b) number of looks) as a function of the number of actual demonstrators for the combined dataset as well as for Exp. 1 and 3 separately. For all depicted results, regression lines were determined with other variables held constant, set to their mean values and the ribbon shows the 95% confidence interval (based on 10 000 simulations of the parameters). Individual datapoints are represented with vertical and horizontal jitter for better visualization. p= 0.1 > + > 0.05 > * > 0.01 > ** > 0.001 > ***

In Exp. 3, we also tested whether the observer pigeons looked specifically at the target location or indistinctly at both the target and distractor location. We found that the first look was significantly more likely directed towards the target location compared to the distractor location in the test condition compared to the control condition (χ2(1) = 5.917, β = 1.005, *p= 0.015*; Fig 2e) and that the interaction term between condition (test, control) and location (target, distractor) significantly predicted the number of looks of the observers (χ2(1) = 5.287, β = -0.644, *p= 0.021*; Fig 2d), indicating that observer pigeons looked more at the target than distractor in the test compared to the control condition. The same trend was apparent for the likelihood of looking although it was not statistically significant (χ2(1) = 3.556, β = -0.565, *p= 0.059*; Fig 2c).

### Collective effect on gaze following

To test the effect of the number of actual demonstrators on the pigeons’ responses, we used the dataset from the test conditions in Exp.1 and 3. In Exp. 1, we found a significant effect of the number of actual demonstrators on the observers’ number of looks (χ2(1) = 6.028, β = 0.583, *p = 0.014*; Fig3b). A similar trend was observed for the observer’s likelihood of looking at the target (χ2(1) = 3.215, β = 0.567, *p = 0.073*; Fig3a, for more details about model results see Suppl. Table S2), although it was not statistically significant. In Exp. 3, we also found a significant effect of number of actual demonstrators on the number of looks of the observers (χ2(1) = 4.535, β = 0.263, *p = 0.033*; Fig3b; the same effect on the likelihood of looking was not significant: χ2(1) = 0.213, β = 0.183, *p = 0.685*; Fig3a). When combining Exp. 1 and 3 into one model, again, we found a significant effect of the number of actual demonstrators on the number of looks (χ2(1) = 8.902, β = 0.324, *p = 0.003*; Fig3b). A similar trend was observed for the likelihood of looking, although not statistically significant (χ2(1) = 3.623, β = 0.285, *p = 0.057*; Fig3a). Furthermore, for all models, the estimated parameters β for the actual number of demonstrator were positive (an increase in the number of actual demonstrator leads to an increase in the observer’s response; see Suppl. Table S2).

To determine whether the configuration of pigeon locations in the test condition— specifically the number of demonstrators present, rather than those actually looking at the target—predicted the observers’ increased response, we replaced the “number of actual demonstrators” variable with the “number of demonstrators” in the model. Overall, this was a weaker predictor of the observers’ response (the likelihood of looking: χ2(1) = 2.464, β = 0.221, *p = 0.117*; the observers’ number of looks; χ2(1) = 3.893, β = 0.207, *p = 0.048*, for more details about model results see Suppl. Table S2), with larger AICs, suggesting a lower performance compared to the original models (likelihood of looking, respectively for the actual number vs. the number of demonstrators: AIC = 685.9, AIC = 690.5; number of looks, respectively: AIC = 1050.9, AIC =1058.4).

To test whether the effect of the number of actual demonstrators on the observers’ responses is linear or nonlinear, we included into the original model a quadratic effect of the number of actual demonstrator for the combined dataset. The quadratic effect was not significant for the likelihood of looking (χ2(1) = 1.504, β = -0.646, *p = 0.220*, for more details about model results see Suppl. Table S2) nor the number of looks (χ2(1) = 1.478, β = -0.556, *p = 0.224*). Additionally, the linear model always had a smaller AIC score than the quadratic model (likelihood of looking, respectively for the linear and the quadratic models: AIC = 685.9, AIC = 697.0; number of looks, respectively: AIC = 1050.9, AIC =1057.0).

## Discussion

Overall, our results supported that 1) pigeons follow the gaze of conspecifics, 2) a larger number of demonstrator pigeons increased the observer pigeons’ gaze following responses, 3) this increase was linear, 4) observer pigeons distinguished the demonstrators’ gaze target, whether it is above or down.

### Gaze following in pigeons

Our results align with previous studies that demonstrate gaze following of conspecifics and human demonstrators across species, including several bird species (ravens: Bugnyar et al., 2004; starlings: Butler & Fernández-Juricic, 2014; geese: Kehmeier et al., 2011; ibises: Loretto et al., 2010; pinguins: Nawroth et al., 2017; ravens: Schloegl et al., 2007; tinamous, emus, rheas and jungle fowls: Zeiträg et al., 2023). However, it is worth noting that the gaze-following responses of our pigeons appeared relatively weak compared to the performance of other bird species in earlier studies (Butler & Fernández-Juricic, 2014; Loretto et al., 2010; Schloegl et al., 2007; Zeiträg et al., 2023), although other studies have reported similar rates (Bugnyar et al., 2004; Kehmeier et al., 2011; Nawroth et al., 2017). The reason for the relatively weak gaze-following responses in our pigeons remains unclear. It may be that pigeons do not follow the gaze of conspecifics as strongly as other species. Alternatively, various contextual factors—such as the pigeons’ experience and subtle environmental influences—may have affected the overall results, warranting further research on gaze following across species.

Moreover, while Experiment 3 demonstrated gaze following in pigeons, Experiments 1 and 2 did not show a significant effect. In Experiment 1, this absence could be explained by the habituation of the demonstrators to the moving objects, leading to weaker observer responses (see Suppl. Mat. 2, Fig S3). Experiment 2, which closely mimicked conventional gaze-following setups from previous studies, ensured that the demonstrator pigeon looked at the target object at least once, but still failed to show clear evidence of gaze following (though some trends in the predicted direction were observed). Experiment 3, however, yielded significant results, likely due to the use of a larger group of new pigeons, which allowed fewer trials and reduced demonstrator habituation, along with improved control conditions.

Interestingly, in Experiment 3, we found that pigeons looked at the target more often than at the distractor, indicating that they specifically oriented their gaze toward the target. This suggests that the observer pigeons could distinguish at least whether the demonstrator pigeons were looking up or down. On these occasions, the demonstrator pigeons typically used one of their laterally projecting foveas to view the target object (92% of the looks towards the target, against 8% for binocular vision). In our table setups, while one fovea was directed at the target (in the direction of the observer pigeons), the other fovea was directed away from the set-up (both target and distractor). Therefore, it is likely that the observer pigeons not only responded to the demonstrators’ head tilts but also specifically followed the gaze of either eye that was directed toward the target.

### Collective effect on gaze following

Our results show that an increased number of demonstrators looking at the target location enhances the observers’ responses. This is largely consistent with human responses in a street context, although we did not observe the saturation effects reported in previous studies (Gallup et al., 2012). To our knowledge, this study is the first to demonstrate a collective effect in non-human animals. Moreover, the increase in observers’ responses is better explained by the number of demonstrators giving a gaze cue toward the target than the number of demonstrators on the other table. This ensured that it was the demonstrators’ looking behavior, in contrast to the spatial configuration of the demonstrators and observers that affected the observers’ gaze following.

The absence of a nonlinear effect in our experiments is likely due to procedural factors, as the maximum number of demonstrators used—nine—was relatively moderate. A saturation effect may still emerge with a larger number of demonstrators. Additionally, as mentioned, the pigeons’ gaze-following response was relatively weak. Nevertheless, our results clearly demonstrated that the collective context significantly influences gaze following in pigeons. This finding is further supported by the fact that our dyadic setup failed to produce evidence of gaze following. One possible explanation is that in group-living species like pigeons, a single individual looking may not be enough to elicit a gaze-following response from others, as the likelihood of any one individual looking at any given moment is high. By contrast, multiple individuals looking in the same direction likely serves as a more reliable indicator of something important in the environment.

An alternative explanation is that an observer is more likely to follow a larger number of demonstrators simply due to increased opportunity; namely, with more demonstrators looking, the observer is more likely to notice at least one gaze cue. However, we designed our table setup so that the observer pigeons could see all demonstrators simultaneously at least with their peripheral vision, with the demonstrators positioned roughly equidistantly from the observers and minimal occlusion between them. Additionally, despite using a dyadic setup and ensuring the demonstrator’s gaze, Experiment 2 failed to reveal a significant effect. This could indicate that one demonstrator is not sufficient to elicit a gaze following response in pigeons.

Overall, we found gaze following in pigeons depends on the number of demonstrators. Quantity or number discrimination is a common cognitive skill present across various species (Agrillo et al., 2007; Bogale et al., 2011; Davis, 1984; Krusche et al., 2010), including pigeons (Emmerton & Renner, 2006, 2009; Watanabe, 1998). Our results suggest that, in addition to other contextual factors—such as dominance in monkeys (Shepherd et al., 2006), gender in humans (Deaner et al., 2006), model species in apes (Hattori et al., 2010; Kano & Call, 2014), facial expressions in both monkeys and humans (Gallup et al., 2014; Goossens et al., 2008), communicative cues in both dogs and humans (Senju & Csibra, 2008; Téglás et al., 2012) —the number of demonstrators may be another factor that modulates gaze following in both humans (Gallup et al., 2012) and pigeons. Further studies are needed to examine this factor in other species.

## Acknowledgements

We would like to thank Drs. Mate Nagy, Dora Biro, Oliver Deussen, and Iain D. Couzin, as well as all the other members of the Max Planck Institute of Animal Behavior and Centre for the Advanced Study of Collective Behaviour, for their insightful feedback and support. Our thanks also go to Mathias Günther for his technical assistance with the use of the facilities. Lastly, we also want to thank Drs. Inge Müller, Daniel Zuniga and the caretakers, for their dedicated care of the pigeons. This study was funded by MPI-AB, the DFG Cluster of Excellence 2117 CASCB (ID: 422037984), and the CASCB BigChunk projects (ID: L21-07), as well as the Grant for Overseas Research by the Division of Graduate Studies, Kyoto University (or DoGS Kyoto University).

## Data and code availability

The data and code used for the analysis can be found on OSF (https://osf.io/bc9gz/)

