## Supplementary material for "Gaze Following in Pigeons Increases with the Number of Demonstrators"

**Suppl Gaze Following**

### Supplementary 1: Dyadic set-up (Exp. 2)

#### Overview of dyadic set-up (Exp. 2)

After Exp. 1, we tested pigeons in a dyadic setup, first using a conspecific (Exp. 2) and then a human demonstrator (Exp. 2 follow-up). The conspecific provided more biologically relevant cues, while the human offered a more controlled, standardized cue. In both experiments, we used a similar tube structure setup as the one used in the group setting (Exp. 1 and 3), modified for the dyad setup. We ran 6 trials for each experiment involving randomized test (demonstrator cue) and control (no demonstrator cue) presentations. However, no significant difference in the observer’s behavior was found between the test and control conditions.

#### Rationale

After conducting the first set of group experiments (Exp. 1), and to show the presence of gaze following in pigeons in a “test vs. control” condition design, we tested our pigeons in a more traditional setup. We ran a few tests in a dyadic setting, where the behavior of a focal subject (observer) was quantified in the presence of one demonstrator giving a gaze cue. We first used a conspecific demonstrator (Exp. 2), then a human demonstrator (Exp. 2 follow-up). Each approach had advantages: whereas a conspecific’s gaze cue was expected to provide more biologically relevant information, a human demonstrator could give a more standardized cue in a more controlled manner. The latter aspect was particularly important, as demonstrator pigeons started to habituate to the moving objects in the tube, and no longer exhibited clear gaze cues in the later trials. It is worth noting that, for Exp. 2, a preliminary attempt failed because the object was too low and the pigeon’s gaze was inadvertently hitting the target, increasing the noise. As a result, this experiment had to be rerun with a higher object. The results for the conspecific dyadic experiment (Exp 2) can be found in the main manuscript, while the human dyadic experiment (Exp. 2 follow-up) is developed hereafter.

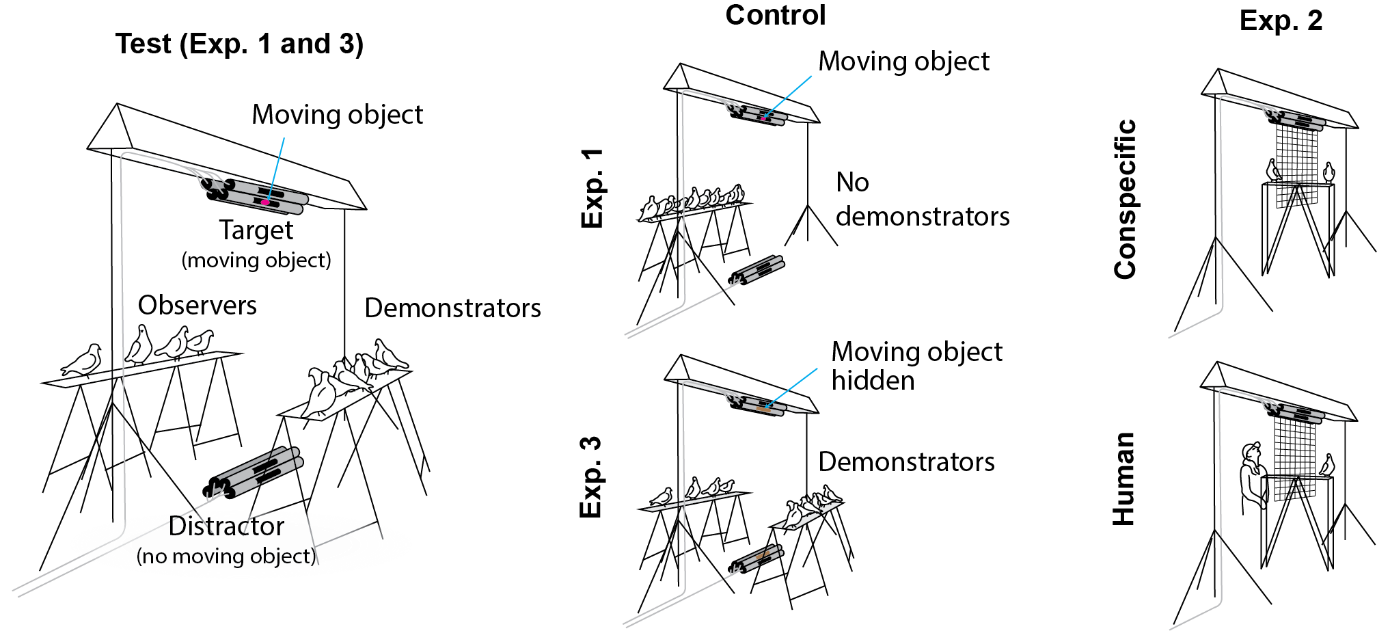

Figure S1: Different setup used in the study. Exp. 1 and 3 were conducted in a group setting, and the test conditions are identical in both, but the control differed. For Exp. 1, all 10 pigeons were moved to the observer side in such a way that there was no demonstrators. For Exp. 3, the windows from some of the tubes were occluded; there were demonstrators but they could not see the moving object and do not give a gaze cue. Exp. 2 was conducted in a dyadic setting, with a conspecific or a human demonstrator.

#### Method (Exp. 2 follow-up)

For the human demonstrator experiment, the bottom object was removed and only the top object was used (Fig S1). One experimenter was standing on one end of the table while the pigeon was placed on the other side of the table (the same table as the dyad conspecific setup). The experimenter and the pigeon were separated by a thin net. The gaze cues were then exhibited in a standard manner (attracting the pigeon’s attention, giving a 5 second gaze cue, looking slightly down for 10 seconds, followed by a 20 seconds break between 2 presentations). The gaze cue was either looking up at the structure (test) or looking behind the pigeon (control). One trial consisted of 3 test and 3 control presentations shuffled in random sequences (each pigeon experiencing 2 trials).

#### Results (Exp 2 follow-up)

We tested if the observers were 1) more likely to look and 2) looking more times at the target location in the presence of a human demonstrator giving a gaze cue (test) compared to not (control). We did not find a significant effect of the condition neither on the likelihood of looking at the target (χ2(1) = 0.15, p= 0.70; see Fig S2 and Table S2), nor the number of looks at the target (χ2(1) = -0.04, p= 1).

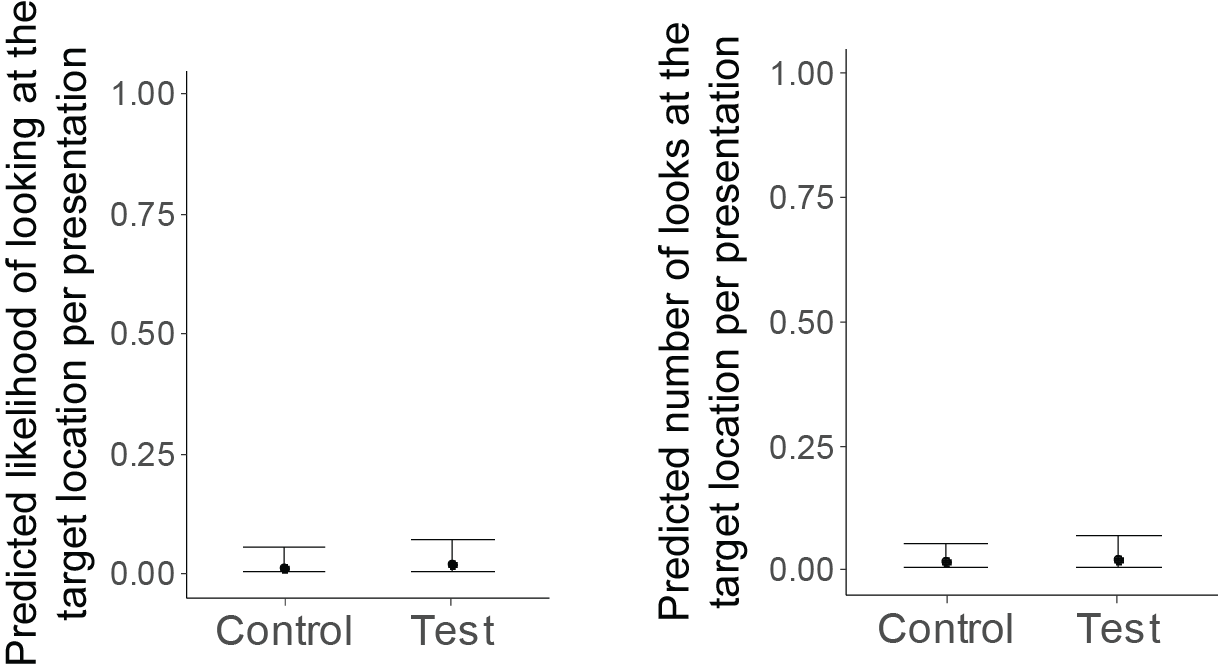

Figure S2: Test-control difference in the likelihood of looking at the target location and the number of looks at the target location for a human demonstrator.

### Supplementary 2: Demonstrators response to the moving objects

To ascertain that the demonstrators’ looking behavior was different in the test and control conditions, we extracted the number of demonstrators that gave a gaze cue in both conditions. In the Test condition of Exp. 1, 69.4% of the demonstrators looked at the target location at least once in the presentation (Fig S3a). Note that for Exp. 1, there was no demonstrator in the control condition, hence the number of “actual” demonstrators cannot be extracted in the control condition. In the dyadic setup (Exp. 2), the demonstrator looked in 97.3% of the test conditions (with a visible moving object) and 36.4% in the control condition (the object is moving but not visible). In Exp. 3, 86.3% of the demonstrators looked at the target object during test presentations against 29.2% during control. This shows that the demonstrators were more likely to look at the object when they could see the moving object compared to when they could not see it. The lower percentage of looks in Exp. 1 compared to Exp. 2 and 3 is likely related to a decrease of the demonstrators' response over the course of the trials related to habituation (Fig S3b).

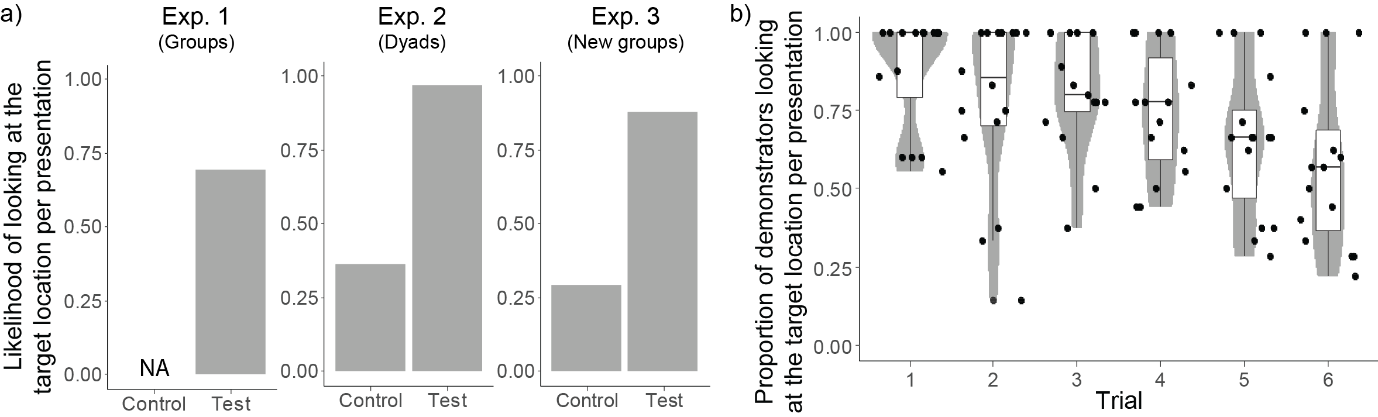
Figure S3: Demonstrators’ response to the moving objects. a) Likelihood of the demonstrators looking at the target location during the test (demonstrators can see the moving object) and control (demonstrators cannot see the moving object) presentations for each experiment. b) Evolution of the proportion of pigeons on the demonstrator table that looked at the object at least once during the presentation over the course of the trials. The distribution is represented using a violin plot (grey shade) and a boxplot (the boxes represent the 0.25 and 0.75 quartiles, with the median represented as a line inside the box, and the whiskers the minimum and maximum values within the lower/upper quartile ± 1.5 times the interquartile range), and the raw observations are displayed with a horizontal jitter for better visualization.

### Supplementary 3: Model details

Table S1: R formulas used for the different models. The models from question 1) were ran with the datasets from Exp. 1, Exp. 2, Exp. 2 follow-up and Exp. 3 (note that for Exp. 2 and Exp. 2 follow-up, the Presentation_location was not included as only the top object was used). The models from question 4) were ran with the dataset from Exp. 3 only as only this dataset showed a significant Test-Control difference. The first 2 models from question 2) were ran with the test data from Exp. 1 and Exp. 3 as well as both datasets combined (note that the Experiment number variable was removed when testing the datasets separately). For the combined dataset, we also tested an alternative model using the Nb_demo (number of demonstrators standing on the table) instead of Nb_actual_demo (number of demonstrators that looked at the target location) to assess the importance of the looking behavior of the demonstrators. Finally, the models from question 3) were ran using the combined dataset to check for the presence of a non-linear effect (saturation).

| Question | | R formula |
| --- | --- | --- |
| Gaze following in pigeons | 1) Comparison test-control | glmer(Looked_or_not ~ Condition + scale(Trial_nb) + scale(Presentation_nb) + scale(Presentation_location) + (Condition + scale(Trial_nb) + scale(Presentation_nb) + scale(Presentation_location)\|Individual_ID), Data, family = binomial(link = "logit")) |
|  |  | glmer(Nb_looks ~ Condition + scale(Trial_nb) + scale(Presentation_nb) + scale(Presentation_location) + (Condition + scale(Trial_nb) + scale(Presentation_nb) + scale(Presentation_location)\|Individual_ID), Data, family = poisson(link = "log")) |
|  | 4) Comparison target-distractor | glmer(Firstlook_is_target ~ Condition + scale(Trial_nb) + scale(Presentation_nb) + scale(Presentation_location) + (Condition + scale(Trial_nb) + scale(Presentation_nb) + scale(Presentation_location)\|Individual_ID), Data, family = binomial(link = "logit")) |
|  |  | glmer(Nb_looks ~ Object * Condition + scale(Trial_nb) + scale(Presentation_nb) + scale(Presentation_location) + (Object * Condition + scale(Trial_nb) + scale(Presentation_nb) + scale(Presentation_location)\|Individual_ID), Data, family = poisson(link = "log")) |
|  |  | glmer(Looked_or_not ~ Object * Condition + scale(Trial_nb) + scale(Presentation_nb) + scale(Presentation_location) + (Object * Condition + scale(Trial_nb) + scale(Presentation_nb) + scale(Presentation_location)\|Individual_ID), Data, family = binomial(link = "logit")) |
| Collective effect | 2) Collective effect | glmer(Looked_or_not ~ scale(Nb_actual_demo) + scale(Experiment) + scale(Trial_nb) + scale(Presentation_nb) + scale(Presentation_location) + (scale(Nb_actual_demo) + scale(Experiment) + scale(Trial_nb) + scale(Presentation_nb) + scale(Presentation_location)\|Individual_ID), Data, family = binomial(link = "logit")) |
|  |  | glmer(Nb_looks ~ scale(Nb_actual_demo) + scale(Experiment) + scale(Trial_nb) + scale(Presentation_nb) + scale(Presentation_location) + (scale(Nb_actual_demo) + scale(Experiment) + scale(Trial_nb) + scale(Presentation_nb) + scale(Presentation_location)\|Individual_ID), Data, family = poisson(link = "log")) |
|  |  | glmer(Looked_or_not ~ scale(Nb_demo) + scale(Experiment) + scale(Trial_nb) + scale(Presentation_nb) + scale(Presentation_location) + (scale(Nb_demo) + scale(Experiment) + scale(Trial_nb) + scale(Presentation_nb) + scale(Presentation_location)\|Individual_ID), Data, family = binomial(link = "logit")) |
|  |  | glmer(Nb_looks ~ scale(Nb_demo) + scale(Experiment) + scale(Trial_nb) + scale(Presentation_nb) + scale(Presentation_location) + (scale(Nb_demo) + scale(Experiment) + scale(Trial_nb) + scale(Presentation_nb) + scale(Presentation_location)\|Individual_ID), Data, family = poisson(link = "log")) |
|  | 3) Non-linear collective effect | glmer(Looked_or_not ~ scale(Nb_actual_demo) + I(scale(Nb_actual_demo^2)) + scale(Experiment) + scale(Trial_nb) + scale(Presentation_nb) + scale(Presentation_location) + (scale(Nb_actual_demo) + I(scale(Nb_actual_demo^2)) + scale(Experiment) + scale(Trial_nb) + scale(Presentation_nb) + scale(Presentation_location)\|Individual_ID), Data, family = binomial(link = "logit")) |
|  |  | glmer(Nb_looks ~ scale(Nb_actual_demo) + I(scale(Nb_actual_demo^2)) + scale(Experiment) + scale(Trial_nb) + scale(Presentation_nb) + scale(Presentation_location) + (scale(Nb_actual_demo) + I(scale(Nb_actual_demo^2)) + scale(Experiment) + scale(Trial_nb) + scale(Presentation_nb) + scale(Presentation_location)\|Individual_ID), Data, family = poisson(link = "log")) |

Table S2: Model results presenting the detailed information for each predictor in every model used in this study, including the beta estimates from the model, as well as the Chi-square value, degrees of freedom, and p-values derived from the likelihood ratio test. Note that the AICs are only comparable for models using the same response, distribution and dataset. Hence, AICs were only provided for models that have at least one other model they can be compared with.

| Question | | Exp. | Response | Type | Variable | # of obs. | Beta | Chi-sq | Df | p | AIC |
| --- | --- | --- | --- | --- | --- | --- | --- | --- | --- | --- | --- |
| Gaze following in pigeons | 1) Comparison test-control | Exp. 1 | likelihood of looking | Test variable | Condition (Test) | 835 | -0.066 | 0.113 | 1 | 0.73631 | - |
|  |  |  |  | Control variable | Trial # | 835 | -0.123 | 1.125 | 1 | 0.28892 |  |
|  |  |  |  |  | Presentation # | 835 | -0.103 | 0.756 | 1 | 0.38461 |  |
|  |  |  |  |  | Presentation location (Top) | 835 | -0.694 | 19.015 | 1 | 0.00001 |  |
|  |  | Exp. 1 | # of looks | Test variable | Condition (Test) | 835 | 0.061 | 0.077 | 1 | 0.78205 | - |
|  |  |  |  | Control variable | Trial # | 835 | -0.050 | 0.179 | 1 | 0.67246 |  |
|  |  |  |  |  | Presentation # | 835 | -0.021 | 0.041 | 1 | 0.83947 |  |
|  |  |  |  |  | Presentation location (Top) | 835 | -0.781 | 20.381 | 1 | 0.00001 |  |
|  |  | Exp 2 | likelihood of looking | Test variable | Condition (Control) | 280 | -0.443 | 1.848 | 1 | 0.17403 | - |
|  |  |  |  | Control variable | Trial # | 280 | -0.030 | 0.035 | 1 | 0.85217 |  |
|  |  |  |  |  | Presentation # | 280 | -0.045 | 0.063 | 1 | 0.80111 |  |
|  |  | Exp 2 | # of looks | Test variable | Condition (Control) | 280 | -0.335 | 2.006 | 1 | 0.15673 | - |
|  |  |  |  | Control variable | Trial # | 280 | -0.141 | 0.901 | 1 | 0.34244 |  |
|  |  |  |  |  | Presentation # | 280 | -0.106 | 0.510 | 1 | 0.47518 |  |
|  |  | Exp 2 (follow-up) | likelihood of looking | Test variable | Condition (Control) | 288 | -0.393 | 0.035 | 1 | 0.85169 | - |
|  |  |  |  | Control variable | Trial # | 288 | 0.003 | -0.042 | 1 | 1 |  |
|  |  |  |  |  | Presentation # | 288 | -0.200 | 0.122 | 1 | 0.72715 |  |
|  |  | Exp 2 (follow-up) | # of looks | Test variable | Condition (Control) | 288 | -0.250 | 0.086 | 1 | 0.76941 | - |
|  |  |  |  | Control variable | Trial # | 288 | 0.214 | 0.264 | 1 | 0.60732 |  |
|  |  |  |  |  | Presentation # | 288 | -0.436 | 1.124 | 1 | 0.28916 |  |
|  |  | Exp. 3 | likelihood of looking | Test variable | Condition (Test) | 670 | 0.448 | 4.964 | 1 | 0.02589 | - |
|  |  |  |  | Control variable | Trial # | 670 | 0.061 | 0.256 | 1 | 0.61268 |  |
|  |  |  |  |  | Presentation # | 670 | 0.032 | 0.081 | 1 | 0.77595 |  |
|  |  |  |  |  | Presentation location (Top) | 670 | -0.445 | 9.497 | 1 | 0.00206 |  |
|  |  | Exp. 3 | # of looks | Test variable | Condition (Test) | 670 | 0.425 | 6.574 | 1 | 0.01035 | - |
|  |  |  |  | Control variable | Trial # | 670 | 0.061 | 0.345 | 1 | 0.55674 |  |
|  |  |  |  |  | Presentation # | 670 | 0.095 | 1.060 | 1 | 0.30316 |  |
|  |  |  |  |  | Presentation location (Top) | 670 | -0.503 | 14.836 | 1 | 0.00012 |  |
|  | 4) Comparison target-distractor | Exp. 3 | First look is target (binary) | Test variable | Condition (Test) | 269 | 1.005 | 5.917 | 1 | 0.01499 | - |
|  |  |  |  | Control variable | Trial # | 269 | -0.075 | 0.074 | 1 | 0.7854 |  |
|  |  |  |  |  | Presentation # | 269 | -0.225 | 0.513 | 1 | 0.47375 |  |
|  |  |  |  |  | Presentation location (Top) | 269 | -1.382 | 20.844 | 1 | < 0.00001 |  |
|  |  | Exp. 3 | # of looks | Test variable | Condition (Test) * Object (Distr) | 1340 | -0.644 | 5.287 | 1 | 0.02149 | - |
|  |  |  |  | Control variable | Trial # | 1340 | -0.055 | 0.527 | 1 | 0.46807 |  |
|  |  |  |  |  | Presentation # | 1340 | 0.042 | 0.396 | 1 | 0.52906 |  |
|  |  |  |  |  | Presentation location (Top) | 1340 | -0.087 | 1.177 | 1 | 0.27788 |  |
|  |  | Exp. 3 | likelihood of looking | Test variable | Condition (Test) * Object (Distr) | 1340 | -0.565 | 3.556 | 1 | 0.05931 | - |
|  |  |  |  | Control variable | Trial # | 1340 | -0.040 | 0.248 | 1 | 0.61834 |  |
|  |  |  |  |  | Presentation # | 1340 | -0.002 | -0.001 | 1 | 1 |  |
|  |  |  |  |  | Presentation location (Top) | 1340 | -0.034 | 0.169 | 1 | 0.68109 |  |
| Collective effect | 2) Collective effect | Exp. 1 | likelihood of looking | Test variable | # of demo that looked | 357 | 0.567 | 3.215 | 1 | 0.07296 | - |
|  |  |  |  | Control variable | Trial # | 357 | -0.453 | 2.572 | 1 | 0.10874 |  |
|  |  |  |  |  | Presentation # | 357 | -0.924 | 13.647 | 1 | 0.00022 |  |
|  |  |  |  |  | Presentation location (Top) | 357 | -0.406 | 2.293 | 1 | 0.12992 |  |
|  |  | Exp. 1 | # of looks | Test variable | # of demo that looked | 357 | 0.583 | 6.028 | 1 | 0.01408 | - |
|  |  |  |  | Control variable | Trial # | 357 | -0.255 | 1.087 | 1 | 0.29723 |  |
|  |  |  |  |  | Presentation # | 357 | -0.728 | 11.742 | 1 | 0.00061 |  |
|  |  |  |  |  | Presentation location (Top) | 357 | -0.409 | 3.332 | 1 | 0.06794 |  |
|  |  | Exp. 3 | likelihood of looking | Test variable | # of demo that looked | 318 | 0.183 | 0.213 | 1 | 0.64481 | - |
|  |  |  |  | Control variable | Trial # | 318 | 0.247 | 1.009 | 1 | 0.31519 |  |
|  |  |  |  |  | Presentation # | 318 | 0.101 | -0.320 | 1 | 1 |  |
|  |  |  |  |  | Presentation location (Top) | 318 | -0.491 | 4.499 | 1 | 0.03391 |  |
|  |  | Exp. 3 | # of looks | Test variable | # of demo that looked | 318 | 0.263 | 4.535 | 1 | 0.03321 | - |
|  |  |  |  | Control variable | Trial # | 318 | 0.226 | 2.507 | 1 | 0.11331 |  |
|  |  |  |  |  | Presentation # | 318 | 0.134 | 1.088 | 1 | 0.29698 |  |
|  |  |  |  |  | Presentation location (Top) | 318 | -0.425 | 6.428 | 1 | 0.01123 |  |
|  |  | Combined (Exp. 1 + 3) | likelihood of looking | Test variable | # of demo that looked | 675 | 0.285 | 3.623 | 1 | 0.05699 | 685.90 |
|  |  |  |  | Control variable | Experiment | 675 | 0.380 | 5.304 | 1 | 0.02128 |  |
|  |  |  |  |  | Trial # | 675 | -0.089 | 0.283 | 1 | 0.59464 |  |
|  |  |  |  |  | Presentation # | 675 | -0.224 | 2.624 | 1 | 0.10524 |  |
|  |  |  |  |  | Presentation location (Top) | 675 | -0.495 | 10.467 | 1 | 0.00122 |  |
|  |  | Combined (Exp. 1 + 3) | # of looks | Test variable | # of demo that looked | 675 | 0.324 | 8.902 | 1 | 0.00285 | 1050.90 |
|  |  |  |  | Control variable | Experiment | 675 | 0.312 | 3.923 | 1 | 0.04762 |  |
|  |  |  |  |  | Trial # | 675 | 0.082 | 0.611 | 1 | 0.43445 |  |
|  |  |  |  |  | Presentation # | 675 | -0.101 | 1.241 | 1 | 0.2652 |  |
|  |  |  |  |  | Presentation location (Top) | 675 | -0.463 | 12.657 | 1 | 0.00037 |  |
|  |  | Combined (Exp. 1 + 3) | likelihood of looking | Test variable | # of demo on the other table | 675 | 0.221 | 2.464 | 1 | 0.11651 | 690.46 |
|  |  |  |  | Control variable | Experiment | 675 | 0.453 | 8.487 | 1 | 0.00358 |  |
|  |  |  |  |  | Trial # | 675 | -0.120 | 0.506 | 1 | 0.47693 |  |
|  |  |  |  |  | Presentation # | 675 | -0.244 | 3.324 | 1 | 0.06827 |  |
|  |  |  |  |  | Presentation location (Top) | 675 | -0.474 | 9.549 | 1 | 0.002 |  |
|  |  | Combined (Exp. 1 + 3) | # of looks | Test variable | # of demo on the other table | 675 | 0.207 | 3.893 | 1 | 0.04848 | 1058.35 |
|  |  |  |  | Control variable | Experiment | 675 | 0.434 | 7.327 | 1 | 0.00679 |  |
|  |  |  |  |  | Trial # | 675 | 0.068 | 0.188 | 1 | 0.66477 |  |
|  |  |  |  |  | Presentation # | 675 | -0.098 | 0.925 | 1 | 0.33626 |  |
|  |  |  |  |  | Presentation location (Top) | 675 | -0.473 | 12.834 | 1 | 0.00034 |  |
|  | 3) Non-linear collective effect | Combined (Exp. 1 + 3) | likelihood of looking | Test variable | # of demo that looked | 675 | 0.925 | 2.683 | 1 | 0.1014 | 696.95 |
|  |  |  |  |  | # of demo that looked squared | 675 | -0.646 | 1.504 | 1 | 0.21999 |  |
|  |  |  |  | Control variable | Experiment | 675 | 0.410 | 6.503 | 1 | 0.01077 |  |
|  |  |  |  |  | Trial # | 675 | -0.103 | 0.373 | 1 | 0.54154 |  |
|  |  |  |  |  | Presentation # | 675 | -0.257 | 3.122 | 1 | 0.07723 |  |
|  |  |  |  |  | Presentation location (Top) | 675 | -0.498 | 10.441 | 1 | 0.00123 |  |
|  |  | Combined (Exp. 1 + 3) | # of looks | Test variable | # of demo that looked | 675 | 0.837 | 3.185 | 1 | 0.07434 | 1057.01 |
|  |  |  |  |  | # of demo that looked squared | 675 | -0.556 | 1.478 | 1 | 0.22415 |  |
|  |  |  |  | Control variable | Experiment | 675 | 0.344 | 4.605 | 1 | 0.03188 |  |
|  |  |  |  |  | Trial # | 675 | 0.056 | 0.134 | 1 | 0.71404 |  |
|  |  |  |  |  | Presentation # | 675 | -0.119 | 1.113 | 1 | 0.29139 |  |
|  |  |  |  |  | Presentation location (Top) | 675 | -0.443 | 12.138 | 1 | 0.00049 |  |
